# KCNG4 genetic variant linked to migraine prevents expression of KCNB1

**DOI:** 10.1101/2024.06.25.600661

**Authors:** Gabriel Lacroix, Shreyas Bhat, Zerghona Shafia, Rikard Blunck

## Abstract

Migraines are a common type of headaches affecting around 15% of the population. The signalling pathways leading to migraines have not been fully understood, but neuronal voltage-gated ion channels have been linked to this pathology, among them KCNG4. KCNG4 (Kv6.4) is a silent member of the superfamily of voltage-gated potassium (Kv) channels, which expresses in heterotetramers with members of the KCNB (Kv2) family. The genetic variant Kv6.4-L360P has previously been linked to migraine, but their mode of action remains unknown. Here, we characterized the molecular characteristics of Kv6.4-L360P when co-expressed with Kv2.1. We found that Kv6.4-L360P almost completely abolishes Kv2 currents, and we propose that this mechanism in the trigeminal system, linked to the initiation of migraine, leads to the pathology.

## Introduction

The predominant role of voltage-gated ion channels is the generation of membrane excitability [3]. The flow of ions through the concerted opening of various ion channels with different selectivity leads to the propagation of the action potentials in neurons. Voltage-gated potassium (Kv) channels, specifically, take the role of repolarization of the membrane following the depolarization caused by the sodium channels [4-6] as well as adjusting excitability of postsynaptic membranes. Kv channels are tetramers where each monomer consists of six transmembrane helices (S1 through S6) [7,8]. S1-S4 form the voltage sensing domain, whose role is to sense the membrane potential. S5-S6 form the pore domain, which lets ions pass through the membrane.

The superfamily of voltage-gated potassium (Kv) channels is divided into 12 families (Kv1-12) [9]. Each Kv family comprises multiple isoforms, which can form functional homo- and heteromers among the family members. The exception are the silent Kv channels Kv5, -6, -8 and -9 [10,11]. The silent Kv channels cannot form functional channels by themselves but require co-expression with members of the Kv2 channel family (Kv2.1-2) to form heteromeric channels altered gating kinetics compared to Kv2 wildtype channels [12-15]. In the case of Kv6, these heteromeric channels comprise alternating subtype (Kv2-6-2-6) [16].

The role of the silent Kv channels is thus to modulate neuronal excitability by co-expression with non-silent channels. While Kv2.1 is ubiquitously expressed, silent subunits have a more restricted expression [14] which would indicate that their role is more tissue-specific by modulating the properties of the membrane in specific positions to produce the required membrane excitability. In the brain, the silent subunits are more abundantly expressed than Kv2 channels, indicating that most Kv2 channels will be regulated by them. Kv6 channels, specifically, are found throughout the brain but not in the retina.

Lafrenière et al. [17] identified the genetic variant KCNG4-L360P in a patient suffering from migraine. Here, we show that Kv6.4-L360P causes almost complete loss of function of Kv2.1 channels when co-expressed in Xenopus oocytes. Xenopus oocytes were selected as a model system because we can control the exact amount of mRNA injected and can exclude any effect during the transcription. Furthermore, Xenopus oocytes do not express Kv2 channels so that we can exclude any influence by endogenous expression.

## Results

Based on a homology model, L360P is located in the middle of the S4-S5 linker connecting the voltage-sensing domain to the pore domain (Fig. 1a-b). The leucine at this position is highly conserved both among different voltage-gated potassium channel families and among different species (Fig. 1 c-d). The dihedral angles of a proline are not compatible with the alpha-helical secondary structure of the S4-S5 linker. The homology model therefore shows a slight kink in the structure but otherwise superposes with the wildtype structure. This is not surprising as it has been generated by Alphafold 3 without further optimization.

**Figure 1.**
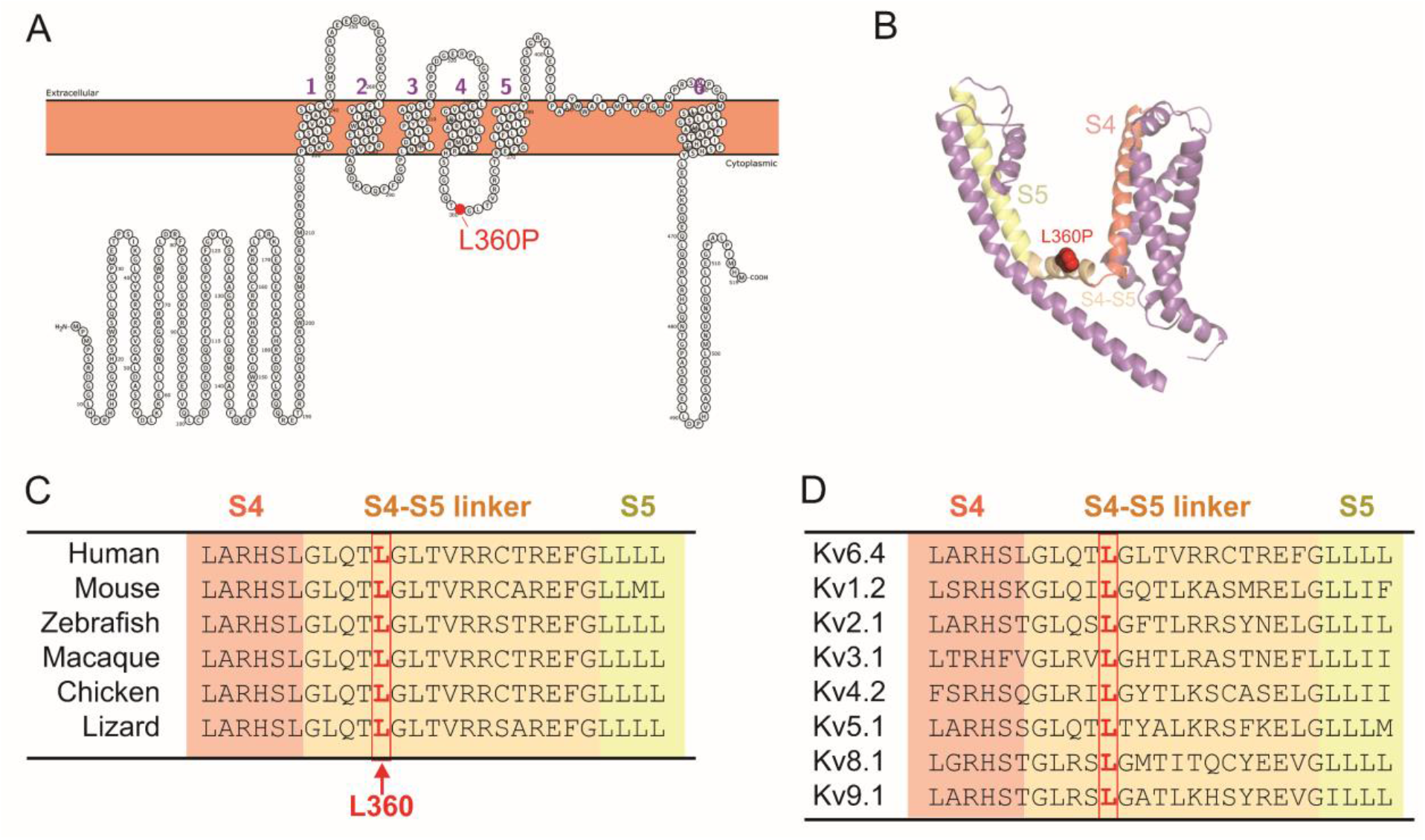
(A) Topology of a Kv6.4 monomer generated by Protter [1]. The mutation L360P is located in the S4-S5 linker. (B) Structural model of the Kv6.4 monomer generated using Alphafold 3 [2]. N- and C-termini are removed for clarity. (C-D) Alignment of the amino acids spanning the end of S4 helix, the S4-S5 linker and the beginning of the S6 helix in Kv6.4 across different species (C) and in different Kv channels (D). The leucine at position 360 in Kv6.4 (highlighted in red) is highly conserved in both instances.

As mentioned above, Kv6.4 channels do not express as homotetramers but have been shown to express in a 2:2 stoichiometry [16]. The Kv2.1/Kv6.4 heteromers show activation and deactivation kinetics very similar to Kv2.1 as reported previously (Fig. 2a). This is reflected in the conductance voltage (GV) characteristics (Fig. 2c), which is only slightly shifted by ∼9 mV and exhibits a shallower slope (Fig. 2c). The apparent gating charge is lowered from z_app_ = 2.37 ± 0.14 to z_app_ = 1.63 ± 0.05. We showed previously [16] that this is caused by the activation being governed by the last subunits to activate, suggesting that Kv6.4 activation is likely shifted to more polarized potentials. In contrast, inactivation, governed by the first subunit to inactivate, is strongly shifted to more polarized potentials (V_½_ = -60.7 mV, Fig. 2b & d). The maximum inactivation in both cases reaches ∼55-60%, given by the ratio of the current of the control pulse before and after the conditioning pulse to different potentials. The apparent gating charge for inactivation is reduced from 4.2 ± 0.2 in Kv2.1 to 3.1 ± 0.1 in Kv2.1/Kv6.4 heteromers.

**Figure 2.**
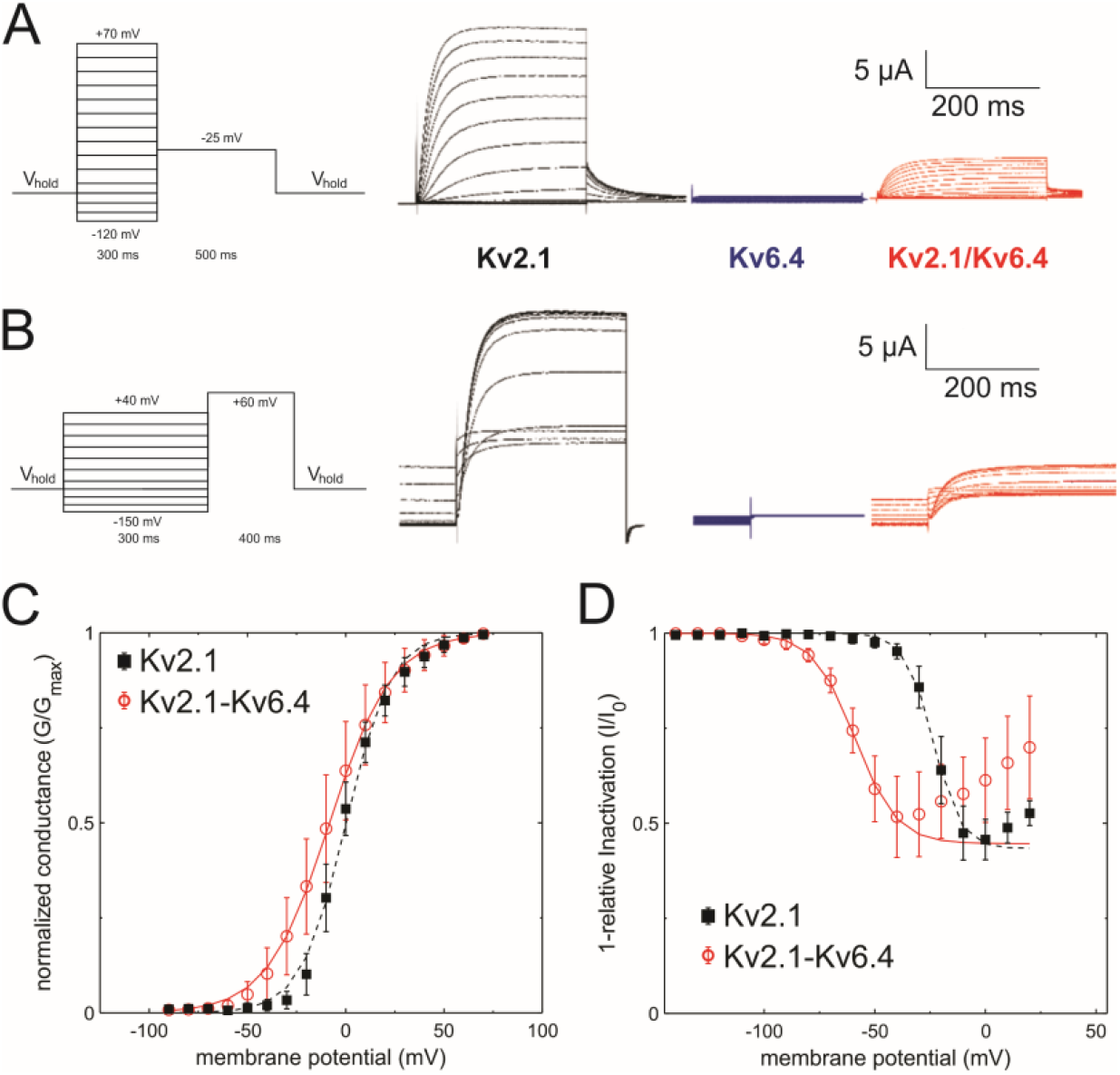
Functional effect of Kv6.4 on Kv2.1. (A) Currents elicited from Xenopus oocytes injected with the indicated constructs in response to membrane potential protocol shown on the left. The conductance protocol begins from the holding potential (−120 mV) to testing potentials in the range from -120 mV to +70 mV in increments of 10 mV, followed by a -25 mV to determine instantaneous GV. (B) Currents elicited from Xenopus oocytes injected with the indicated constructs in response to membrane potential protocol shown on the left. The inactivation protocol begins at a holding potential (−120 mV) to a range of 4 seconds long inactivating pulses in the range of -150 mV to +40 mV in increments of 10 mV, followed by a depolarizing pulse to +60 mV in order to measure the level of inactivation caused by the preconditioning pulse (C) Conductance-voltage curve for Kv 2.1 (N=8) and Kv2.1/Kv6.4 (N=18). The curves were fitted with a single Boltzmann equation. (D) Inactivation-voltage curve for Kv2.1 (N=9) Kv2.1/Kv6.4 (N=11). These curves were fitted with a single reverse Boltzmann equation.

To estimate the effect of the migraine-linked genetic variant Kv6.4-L360P, we generated the mutation in Kv6.4-cDNA and in vitro transcribed them before expressing them in Xenopus oocytes, that allow us proper control of the relative expression level of Kv2.1 and Kv6.4 constructs. We expressed Kv6.4-L360P alone and co-expressed it with Kv2.1 in a 10:1 (w 6.4-L360P : w 2.1) ratio. A 10-fold excess of the silent subunit ensures that the expression of Kv2.1-homotetramers background expression is reduced to a minimum. As expected, no functional expression was observed in the absence of Kv2.1 (Fig. 3a). Co-expression with Kv2.1 led to very small currents after three days of expression, compared to 1 day for Kv6.4 wildtype. There was no significant difference discernable between the currents, kinetics and voltage dependencies compared to the Kv2.1 homotetramers (Fig. 3a-b, Table 1). The delayed and low expression and the fact that the characteristics resembled Kv2.1 alone suggested that we are merely recording Kv2.1 homotetrameric background expression. Accordingly, the currents were small and displayed only after longer incubation time (18-24 vs 66-72 hrs; Table 2).

**Figure 3.**
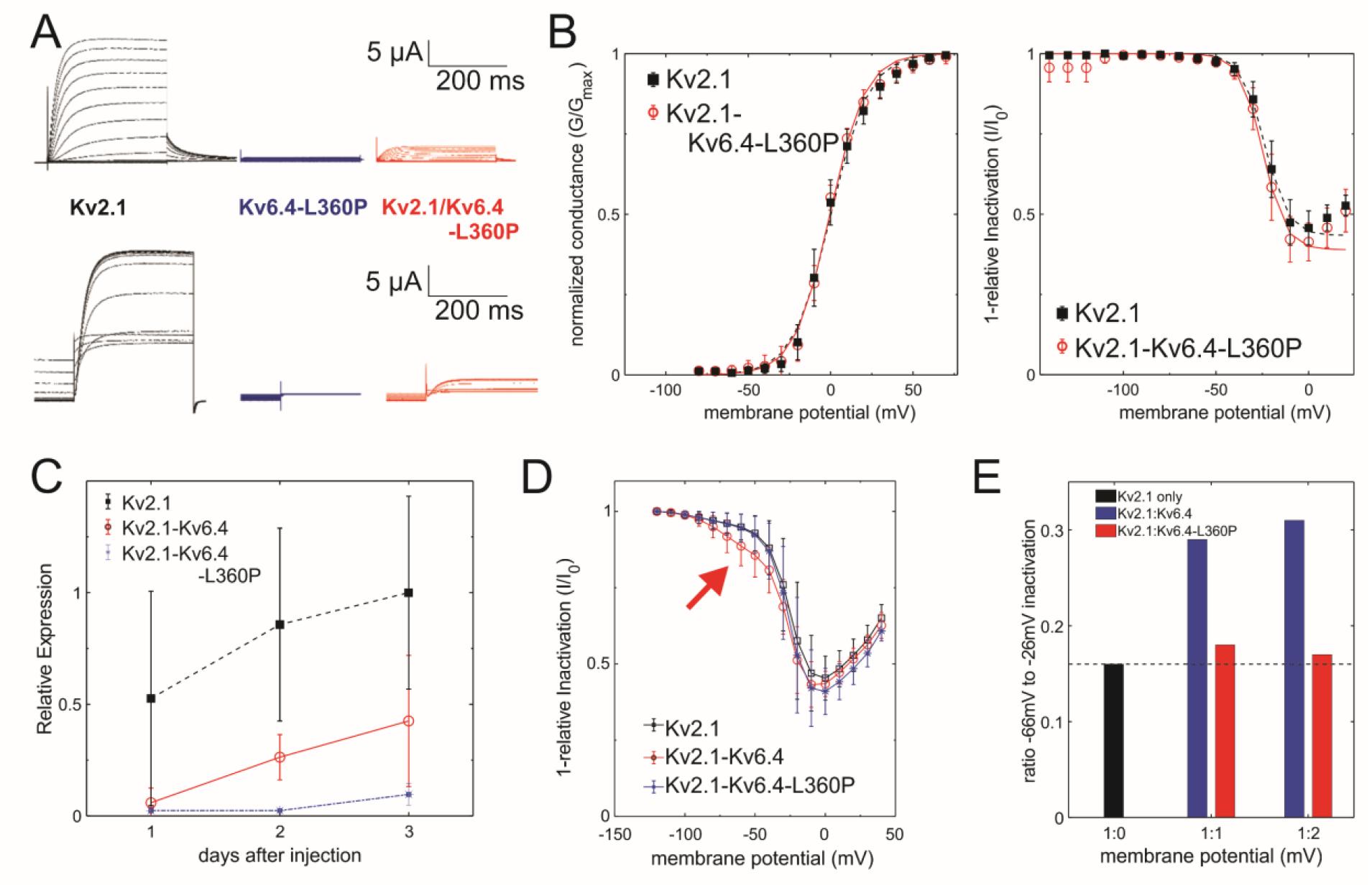
(A) Currents elicited from elicited from Xenopus oocytes injected with the indicated constructs in response to membrane potential protocols shown in Fig. 1a-b. (B) left: Conductance-voltage curve for Kv2.1 (N=8), Kv2.1/Kv6.4 (N=18) and Kv2.1/Kv6.4-L360P (N=9). Right: Inactivation-voltage curve for Kv2.1 (N=9), Kv2.1/Kv6.4 (N=11) and Kv2.1/6.4-L360P (N=6). All curves were fitted with a single Boltzmann equation. Fitting results are summarized in Table 1. (C) Intensity of currents measured obtained after different incubation times for Kv2.1 (N=36), Kv2.1/Kv6.4 (N=31) and Kv2.1/6.4-L360P (N=49). (D) Conductance-inactivation curves for 1 ng Kv2.1 and variably 1 ng Kv6.4 or Kv6.4-L360P. (E) Recordings in D in 1:1 and 1:2 ratio were fit to a superposition of two Boltzmann distributions. The fraction of the inactivation curve with V_50_ = -66 mV is shown.

**Table 1.** Results from a single Boltzman fit for the Kv2.1, Kv2.1/Kv6.4 and Kv2.1/Kv6.4-L360P.

| **Conductance** | **V50 (mV)** | **Slope (mV)** | **Apparent charge** | **N** |
| --- | --- | --- | --- | --- |
| Kv2.1 | $0.1 \pm 0.8$ | $10.9 \pm 0.7$ | $2.37 \pm 0.14$ | 6 |
| Kv2.1 / Kv6.4 | $-9.1 \pm 0.6$ | $15.9 \pm 0.5$ | $1.63 \pm 0.05$ | 18 |
| Kv2.1 / Kv6.4-L360P | $-0.41 \pm 0.63$ | $10.7 \pm 0.6$ | $2.41 \pm 0.12$ | 9 |

| **Inactivation** | **V50 (mV)** | **Slope (mV)** | **Apparent charge** | **Maximum Inactivation** | **N** |
| --- | --- | --- | --- | --- | --- |
| Kv2.1 | $-23.8 \pm 0.37$ | $-6.0 \pm 0.3$ | $-4.19 \pm 0.21$ | $0.58 \pm 0.02$ | 9 |
| Kv2.1 / Kv6.4 | $-60.7 \pm 0.56$ | $-8.4 \pm 0.4$ | $-3.07 \pm 0.14$ | $0.55 \pm 0.02$ | 11 |
| Kv2.1 / Kv6.4-L360P | $-24.7 \pm 0.61$ | $-6.1 \pm 0.5$ | $-4.24 \pm 0.33$ | $0.61 \pm 0.02$ | 6 |

**Table 2.**
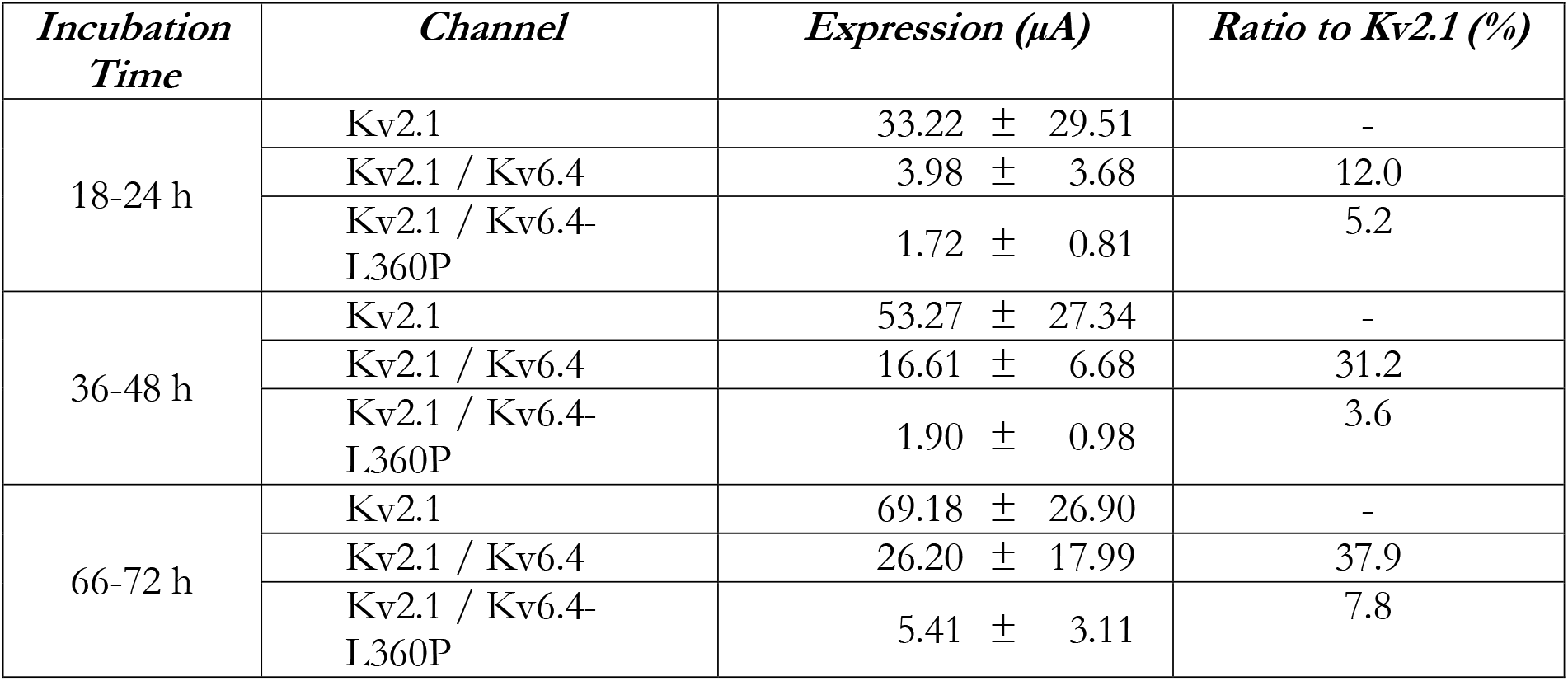
Level of expressions for Kv2.1 (N=36), Kv2.1/Kv6.4 (N=31) and Kv2.1/Kv6.4-L360P (N=49). The oocytes were injected with 5ng of Kv2.1 and 50ng of Kv6.4 or Kv6.4-L360P depending on the channel studied. For incubation times 18-24 h and 36-48 h with Kv2.1 / Kv6.4-L360P, most of the currents are leak. Only 66-72 h had usable currents.

To ensure that the absence of channels is not due to too slow recovery from inactivation, we tried to apply longer and more hyperpolarized prepulses, which had no effect. While unlikely, it would be possible that the mutation L360P in Kv6.4 removes the strongly shifted inactivation in Kv6.4. It was recently suggested that a leucine in the S6 of Kv2.1 that interacts with another leucine in the S4-S5 linker is responsible for inactivation in Kv2.1 channels [18]. If this were the case, however, we would not observe a reduction in the expression (Fig. 3c).

The increase in homotetrameric Kv2.1 over time, compared to Kv2.1-Kv6.4 heteromers while expressing delayed with respect to Kv2.1 homotetramers (Fig. 3c) suggests that Kv2.1 is retained longer in the plasma membrane. We next investigated the possibility that Kv6.4-L360P is expressing much stronger than Kv6.4 wildtype. In a previous study, we observed that Kv2.1-Kv6.4 complexes in a 3:1 ratio were produced but remained in the ER and were not trafficked to the plasma membrane [16]. We therefore increased the proportion of Kv2.1 in the expression to 1:2 and 1:1 (Kv2:Kv6). For Kv6.4 wildtype, the inactivation characteristics showed two distinct populations following the Kv2.1-Kv6.4 heteromers (V_50_ = -66 mV) and the Kv2.1 homotetramers (V_50_ = -26 mV; Fig. 3d, *red arrow*). This component accounts for approximately 30% of the inactivation (Fig. 3e). For Kv6.4-L360P, however, not even a small fraction of the shifted inactivation curve of Kv2.1-Kv6.4 heteromers was observed. The channels behaved identically to Kv2.1 homotetramers, indicating that Kv6.4-L360P abolishes expression of any functional Kv2-Kv6 complexes.

### Kv2.1/Kv6.4-L360P complexes are strongly produced

Considering that the expression of Kv2.1 is strongly reduced in the presence of the Kv6.4-L360P mutant, the mutation cannot simply lead to a misfolded protein, but Kv2.1-Kv6.4-L360P complexes have to get assembled. To verify this, we undertook immunoblotting experiments. We used Kv6.4 fused to enhanced GFP (Kv6.4-eGFP) and co-expressed the wildtype and L360P variant with Kv2.1 wildtype as well as on its own. Expression was detected via the eGFP fluorescence from the acrylamide gel. Both Kv6.4 and Kv6.4-L360P were expressed in absence and presence of Kv2.1 in approximately equal amounts. However, in both cases, expression relative to the Aquaporin 2-RFP control was significantly higher for Kv6.4-L360P.

We then tested expression in a mammalian expression system; Kv2.1 and Kv6.4-L360P were co-expressed in HEK293 cells (Fig. 4a). As expected, Kv2.1 alone is highly expressed, and expression is significantly lowered in the presence of Kv6.4 wildtype (Fig. 4b). Increasing the amount of Kv6.4 relative to Kv2.1 further decreased expression. The expression is not lowered, however, in the presence of Kv6.4-L360P compared to Kv6.4-WT, indicating that Kv6.4-L360P is translated and folded properly and forms heterotetramers with Kv2.1. These channels are either not trafficked to the plasma membrane or are not functional.

**Figure 4.**
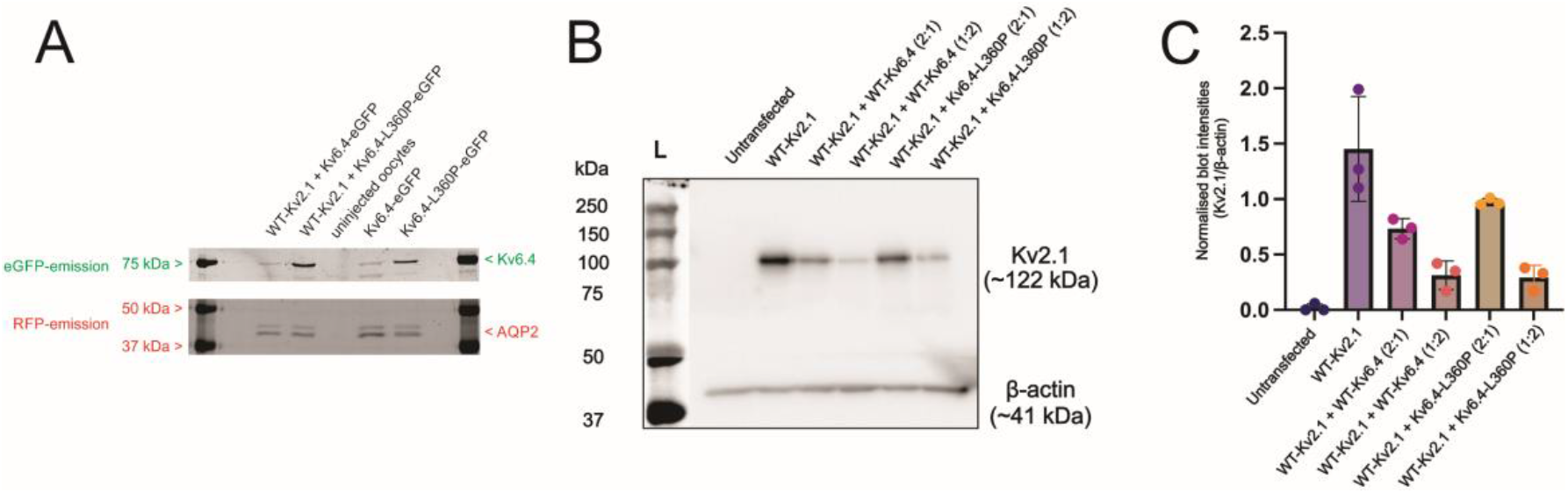
(A) SDS page gel from full membranes obtained from Xenopus oocytes. Kv6.4 is fused with eGFP and AQP2 with RFP for detection. eGFP and RFP fluorescence were obtained using appropriate filtersets. The same gels were imaged with both filtersets and superposed here. Aquaporin shows a double band due to glycosylation (N=3). (B) HEK293 cells, on confluency, were transiently transfected with either WT-Kv2.1 (2 μg), WT-Kv2.1:WT-Kv6.4 (2 μg: 1 μg), WT-Kv2.1:WT-Kv6.4 (2 μg: 4 μg), WT-Kv2.1:Kv6.4-L360P (2 μg: 1 μg) or WT-Kv2.1:Kv6.4-L360P (2 μg: 4 μg). A total of 6 μg of plasmid DNA were transfected each condition topped with pcDNA3.1 as a control vector. Untransfected cells were taken as negative controls (first lane). Membrane proteins extracted from these cells were denatured and resolved with SDS-PAGE. After transfer of the proteins onto nitrocellulose membranes, the blots were incubated overnight at 4 °C with anti-Kv2.1 (top) or anti-β-actin (bottom, loading control) antibody. The immunoreactive bands were detected with fluorescently labeled secondary antibody. The blot is representative of three independent experiments. (C) Densitometric intensities of protein bands of Kv2.1 were normalized to those of β-actin in each condition and the resulting ratios were plotted as box plot that show mean values ± S.D.

## Discussion

The role of silent Kv channels (KvS) is to modulate Kv2 currents, primarily reflected in the voltage dependence of inactivation, shifting its voltage-dependence towards more polarized potentials compared to the Kv2 homomer (Fig. 2d). While the Kv2 subunits are more ubiquitously expressed throughout the nervous system, the differential expression patterns of the silent subunits leads to a spatial distribution of the functional phenotypes of the Kv2-KvS complexes. The KvS thus play a regulatory role reducing the availability of Kv2 by (i) lower expression and (ii) increased subthreshold inactivation.

The mutation Kv6.4-L360P characterized here is located in the center of the S4-S5 linker. The S4-S5 linker region is essential in Kv for the coupling between voltage-sensor domain, which responds to changes in the membrane potential, and pore domain, which acts as effector opening the ion conduction pathway. The coupling is accomplished by annealing to the S6 C-terminus. This link allows for the movement of the S4 to be transmitted to the pore and lets the channel open when the voltage sensors are activated. Mutations in this region in Shaker Kv channels [19] modulate coupling between the voltage sensor and the pore leading to a shifted voltage dependence for the channel, while simultaneously the S4 voltage sensor becomes more sensitive to the membrane potential. The change from a leucine to a proline means a reduction in size and in hydrophobicity and disturbs the alpha-helix forming the S4-S5 linker [20]. Given the position of the mutation, one might suspect the channels to be expressed but not opening due to disturbed coupling. In this case, however, we should have observed gating currents, as observed for uncoupling mutants in Shaker Kv channels [19]. But gating currents were not observed.

In Kv2.1 the analogous position L316 has been suggested to be involved in the development of inactivation. The mutant L316A led to accelerated inactivation and the inactivation-voltage relation was shifted to more polarized potentials by interaction with the S6 [18]. This would allow for the possibility that the L360P mutation alleviates the strong shift of the Kv6.4-inactivation although it would still be surprising to find the exact value as in Kv2.1. Two other observations argue against this interpretation. First, we observed a strong reduction in functional channels (Figs. 2-3) despite the fact that Kv2.1 and Kv6.4-L360P are translated as well as Kv2.1 alone (Fig. 4), indicating that the assembled channels are either not functional or not trafficked. Secondly, we recently studied 10 different genetic variants in KCNB2 (Kv2.2) linked to neurodevelopmental disease [21]. These genetic variants were spread throughout the channel in the 3 dimensional fold but still all influenced inactivation in KCNB2, indicating that not only the coupling region has a strong influence on the development of inactivation.

Many genetic variants in genes encoding for Kv channels are linked to a variety of neurological and cardiac diseases. The silent Kv, specifically, were linked to cone dystrophy in the eye leading to a loss of sight (Kv8.2 coexpressed with Kv2.1)[22] and epilepsy (Kv8) [23,24]. Kv6.4 has not only been linked to migraine in the variant characterized in the present study [17] but also to reduced labor pain when expressed in uterine sensory neurons. The variant Kv6.4-V419M were retained in the ER and prevented Kv2.1 from reaching the membrane. This leads to higher excitation threshold but longer interspike interval in the neurons, thereby reducing labor pain [25].

The mutation L360P in Kv6.4 likely has a similar effect. Migraine has been suggested to develop in the trigeminal ganglion [26,27]. Both Kv2.1 and Kv6.4 express in the trigeminal ganglion, specifically in the neurofilaments [28-30]. As in the uterine sensory neurons, not all neurofilaments are positive for Kv6.4 expression [27]. We can speculate that the downregulation of Kv2.1 in the neurofilaments of the trigeminal ganglion by co-expression with Kv6.4-L360P likely follows a similar mechanism as the V419M variant in the uterine sensory neurons and thereby causes migraine.

## Materials and Methods

### Pulse Protocols and Data Analysis

Voltage protocols for the activation and inactivation measurements are shown in figure 1. Holding potential was adjusted based on the potential at which the measured channels opened. The normalized activation curve was fitted to a Boltzmann equation 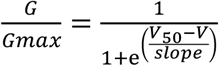 where V_50_ is the voltage at which 50% of the channels are open or inactivated and *slope* the slope factor. The apparent charge *z* was defined as 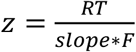 where *T* is the temperature in Kelvins (298 K in our case) and *R* is the ideal gas constant 8.314 J mol^-1^K^-1^. The inactivation was fitted with a reverse Boltzmann equation 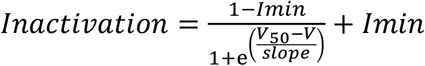 where V50 is the voltage at which 50% of the channels are open or inactivated, *slope* the slope factor. *I*_*min*_ is the lowest point of the curve, usually around 0.4 meaning 60% inactivation.

### Expression of Kv2/Kv6 in Xenopus oocytes

Kv2.1 and Kv6.4–eGFP were introduced into the pBSta and pSP64 vector, respectively, and cRNA was in vitro transcribed by using a T7 mMachine kit (Invitrogen), according to the manufacturer’s protocol. The L360P mutation was introduced in Kv6.4-eGFP by PCR mutagenesis and verified by sequencing. Oocytes from *Xenopus laevis* were surgically obtained, according to protocols approved by the Comité de déontologie de l’expérimentation sur les animaux de l’Université de Montréal. Oocytes were injected with mRNA (1) 1 ng Kv2.1, (2) 5 ng Kv2.1 + 50 ng Kv6.4–eGFP and (3) 5 ng Kv2.1 + 50 ng Kv6.4-L360P–eGFP. Injected oocytes were incubated for 24 to 48 h at 18 °C.

### Immunoblotting of Kv2.1 and Kv6.4

#### Xenopus oocytes

Per condition, 10 injected oocytes were homogenized in 200μl phosphate buffered saline (PBS), filled up to 1,000 μl and centrifuged 5 minutes at 180 x *g*. The supernatant was collected and centrifuged for 20 minutes at 20,000 x *g*. The pellet was collected and resuspended in 30 μl PBS and 10 μl Laemmli buffer was added. 16 μl were loaded on a 10% acrylamide gel.

The samples were injected with mRNA (1) 5 ng Kv2.1 and 50 ng Kv6.4–eGFP, (2) 5 ng Kv2.1 and 50 ng Kv6.4-L360P-eGFP, (3) 50 ng Kv6.4-eGFP, (4) 50 ng Kv6.4-L360P-eGFP as well as (5) uninjected oocytes. Every injected oocyte was co-injected with 5 ng of AQP2-RFP for loading control. The calibration ladder used was Precision Plus Protein Dual Color Standards (BioRad, #1610374). The same gel was imaged with eGFP and RFP filtersets, i.e. blue excitation light and green imaging filter for eGFP and green excitation light and red imaging filter for RFP.

#### Mammalian Cell Lines

HEK293 cells were seeded in Dulbecco’s modified Eagle’s medium containing 10% fetal calf serum and 1% pencillin/streptomycin on day 1 at a density of 300,000 cells/well in 6 well plates. On day 2, each well was transfected with a fixed concentration of Kv2.1 plasmids (2 μg) and variable concentrations of either WT-Kv6.4 or Kv6.4-L360P (0, 1 or 4 μg) using an in-house polyethylene imine transfection reagent. Total amount of DNA transfected per well was bought up to 6 μg using pcDNA3.1 as a control vector. On day 3, the cells were washed 3 times with ice-cold phosphate-buffered saline and lysed with a buffer containing Tris·HCl, pH 8.0, 150 mm NaCl, 1% dodecyl maltoside, 1 mm EDTA, and protease inhibitors (CompleteTM, Roche Applied Science). The detergent-insoluble material was subsequently removed by centrifugation (16,000 × g for 15 min at 4°C) and the protein concentration of the supernatant was determined. 20 μg of total proteins per condition were mixed with sample buffer containing SDS and β-mercaptoethanol, denatured at 45°C for 30 min, and electrophoretically resolved in denaturing polyacrylamide gels. After transfer of the proteins onto nitrocellulose membranes, the blots were probed with an antibody against either Kv2.1 (rabbit, ab194523) or β-actin (mouse, ab49900) at recommended dilutions. This was followed by the addition of horseradish peroxidase (HRP)-conjugated secondary antibody (1:3000) and the ensuing immunoreactivity was detected by chemiluminescence and quantified using ImageJ.

## Data Availability

All data and materials will be made available upon request.

## Acknowledgements

The authors would like to thank Yoann Lussier for technical assistance. This work was financially supported by the Canadian Institutes of Health Research (CIHR, PJT-169160) and the Natural Sciences and Engineering Research Council of Canada (NSERC, RGPIN-2023-04752). CIRCA is a research center financially supported by the Fonds de recherche Québec — Santé.

